# Structurally distinct oligomers of islet amyloid polypeptide mediate toxic and non-toxic membrane poration

**DOI:** 10.1101/095158

**Authors:** Melissa Birol, Sunil Kumar, Elizabeth Rhoades, Andrew D. Miranker

## Abstract

Peptide mediated gain-of-toxic function is central to pathology in Alzheimer’s, Parkinson’s and diabetes. In each system, self-assembly into oligomers is observed and can also result in poration of artificial membranes. Structural requirements for poration and the relationship of structure to cytotoxicity is unaddressed. Here, we focus on islet amyloid polypeptide (IAPP) mediated loss of insulin secreting cells in diabetics. Newly developed methods enable structure-function inquiry to focus on intracellular oligomers composed of hundreds of IAPP. The key insights are that porating oligomers are internally dynamic, grow in discrete steps and are not canonical amyloid. Moreover, two class of pores coexist; an IAPP-specific ligand establishes that only one is cytotoxic. Toxic rescue occurs by stabilizing non-toxic poration without displacing IAPP from mitochondria. These insights illuminate cytotoxic mechanism in diabetes and also provide a generalizable approach for inquiry applicable to other partially ordered protein assemblies.

**Highlights:**

- The peptide amyloid precursor, IAPP, forms two classes of membrane porating oligomers.
- The two classes have a >100-fold difference in pore size with the large pore form correlated with mitochondrial depolarization and toxicity.
- A drug-like molecule distinguishes between the two oligomer classes and rescues toxicity by stabilizing non-toxic poration without displacing IAPP from the mitochondria.
- The mechanism of pore-forming oligomer assembly includes stepwise coalescence of smaller, dynamic assemblies.

## Introduction

Achieving molecular scale insights in structure-function faces unique challenges when function arises from oligomers formed from partially ordered proteins. Attention to proteins in this class has dramatically increased in recent years as such systems are increasingly being found to have functional roles in biology (Courchaine et al., 2016). Dynamic assemblies can occur in aqueous and membrane milieus, for example, giving rise to nucleoli (Brangwynne et al., 2011) and signaling complexes (Su et al., 2016) respectively. As is often the case in biology, functional observations such as these are a product of selective pressure optimizing an intrinsic physical property of polypeptides. Unchecked, however, this property can also result in the uncontrolled assembly of normally monodisperse proteins. These entities have gains-of-function the downstream consequences of which are cytotoxicity and/or irreversible fiber formation (Aguzzi and Altmeyer, 2016).

Pathological self-assembly of proteins underpins numerous important human diseases including Alzheimer’s, Parkinson’s and type II diabetes (Knowles et al., 2014). In each case, a different protein and tissue is involved; however, the broad characteristics include a fibrous end-product termed amyloid. Amyloid fibers are defined by an organization in which β-strands are arranged orthogonal to the filament axis. Although amyloid fibers are a hallmark, it has been widely shown that pre-amyloid assemblies are the dominant source of toxic gains-of-function. Pre-amyloid toxins can occur spontaneously in solution, by encounter with a biological polyanion such as a phospholipid bilayer, nucleic acid or recently, polyphosphate (Cremers et al., 2016) and by templated assembly on the lateral walls of preexisting fiber (Knowles et al., 2014; Schlamadinger and Miranker, 2014).

Islet amyloid polypeptide (IAPP) is a 37-residue peptide-hormone copackaged and cosecreted with insulin by the endocrine pancreas. In type II diabetics, this peptide is observed to form amyloid fibers that are associated with loss of β-cell mass (Mukherjee A, 2015). Orthologues and mutational studies reveal a strong correlation between amyloid formation potential and cytotoxicity. Nevertheless, it is the structurally poorly-defined, oligomeric species that are present prior to fibrillar aggregates that appear cytotoxic. In solution, IAPP weakly samples a-helical conformations that are stabilized upon interaction with phospholipid bilayers (Williamson et al., 2009). Membrane catalyzed self-assembly follows binding with gains-of-function that include energy independent membrane translocation, membrane poration and mitochondrial localization (Costes et al., 2013). At the molecular level, the relationships of structures to functions, including downstream observations of mitochondrial dysfunction and cell death, remain unclear.

Membrane interacting oligomers whose structures and dynamics result in poration are widely believed to underpin IAPP’s toxic gain-of-function (Mukherjee et al., 2015). Numerous studies using synthetic vesicles have provided evidence for a plethora of alternative poration mechanisms: discrete protein pore formation, membrane disruption through carpeting, removal of phospholipid like a surfactant or as response to the induction of surface tension (Last et al., 2013). Which, if any, of these mechanisms is relevant to cells is unknown, in part because many basic properties of poration, such as pore size, remain unaddressed. The role of oligomer structure in these phenomena is also unclear although there is indirect evidence, such as the near loss of poration upon truncation of the non-membrane bound and disordered C-terminal residues (Brender et al., 2012). Additionally, as IAPP localizes to mitochondria in cultured cells in a manner correlated with cell death, it is an appealing, although unproven idea that IAPP directly mediates poration of mitochondria (Kegulian et al., 2015). In studies unrelated to IAPP, small molecule mediated poration of mitochondria, using 2, 4-Dinitrophenol (DNP), has even been shown to increase the viability of β-cells and rescue metabolic syndrome in animal models (Perry et al., 2015). How does one reconcile these conflicting observations? Clearly a deeper insight is required to understand the molecular basis of IAPP mediated poration and its relationship to toxicity.

To address this challenge, we carried out parallel measurements of IAPP on membrane models and inside live cells. Synergy between cellular and model measurements was achieved, in part, by engineering a permeation assay that uses a minimally processed, cell derived membrane. Using this assay, two sequentially sampled forms of poration are newly identified. A previously developed IAPP-specific ligand, ADM-116, is used here as a tool that shows toxic mitochondrial depolarization to be associated with only one of the two forms. A structure-based fluorescence approach, which we term diluted-FRET, is developed that brings focus onto large, homooligomeric assemblies of IAPP. We find that functional oligomers contain scores to hundreds of IAPP, populate several discrete states and none of these states is canonical amyloid. Our results support a model whereby IAPP-mediated toxicity derives from large, mitochondrial membrane associated assemblies of IAPP.

## Results

### Leakage profiles using a cell derived membrane

IAPP induces leakage in giant plasma membrane vesicles (GPMVs). GPMVs were prepared from live INS-1 cells as previously described (Schlamadinger and Miranker, 2014; Sezgin et al., 2012) with an additional step of pre-staining cytoplasmic components with a thiol-reactive fluorescent probe, CellTracker Orange. These GPMVs are stable for days with little loss of fluorescence even under prolonged confocal imaging (Figure 1A). Membrane leakage kinetics of 5 μM GPMVs (expressed as apparent concentration in phospholipid monomer units) were monitored as a loss in fluorescence intensity after co-addition of IAPP and 1 mM of the small molecule fluorescent quencher, acid-blue 9 (AB9) (Figure 1D). Loss of intensity in the presence of a quencher comes both from material loss from the lumen as well as influx of the quencher into the GPMV. At 2:1, 5:1 and 10:1, protein:lipid (P:L), single exponential decay is observed, ending in complete loss of fluorescence (Figure 1D). Rate constants derived from fits are 42 ± 6 s^-1^, 54 ± 5 s^-1^ and 60 ± 3 s^-1^ observed for 2:1, 5:1, and 10:1 P:L, respectively. This suggests that holes large enough for the small molecule quencher are generated with an efficiency that is only weakly dependent of protein concentration. Note that the high P:L ratios used here are typically associated with destruction of synthetic lipid vesicles (Kegulian et al., 2015). However, these P:L are consistent with standard application of amyloid precursors to cell culture where the addition of low μM protein to a culture plate corresponds to P:Ls of 100:1 or more. This routine, 5+ order of magnitude disparity, has been noted as a potential cause for concern when interpreting solution biophysical results (Wimley, 2010). That we observe no gross changes to the GPMV morphology at these P:L suggests that our model membrane retains the robustness of the plasma membrane, possibly a consequence of the retention of some cytoskeletal components (Podkalicka et al., 2015).

**Figure 1:**
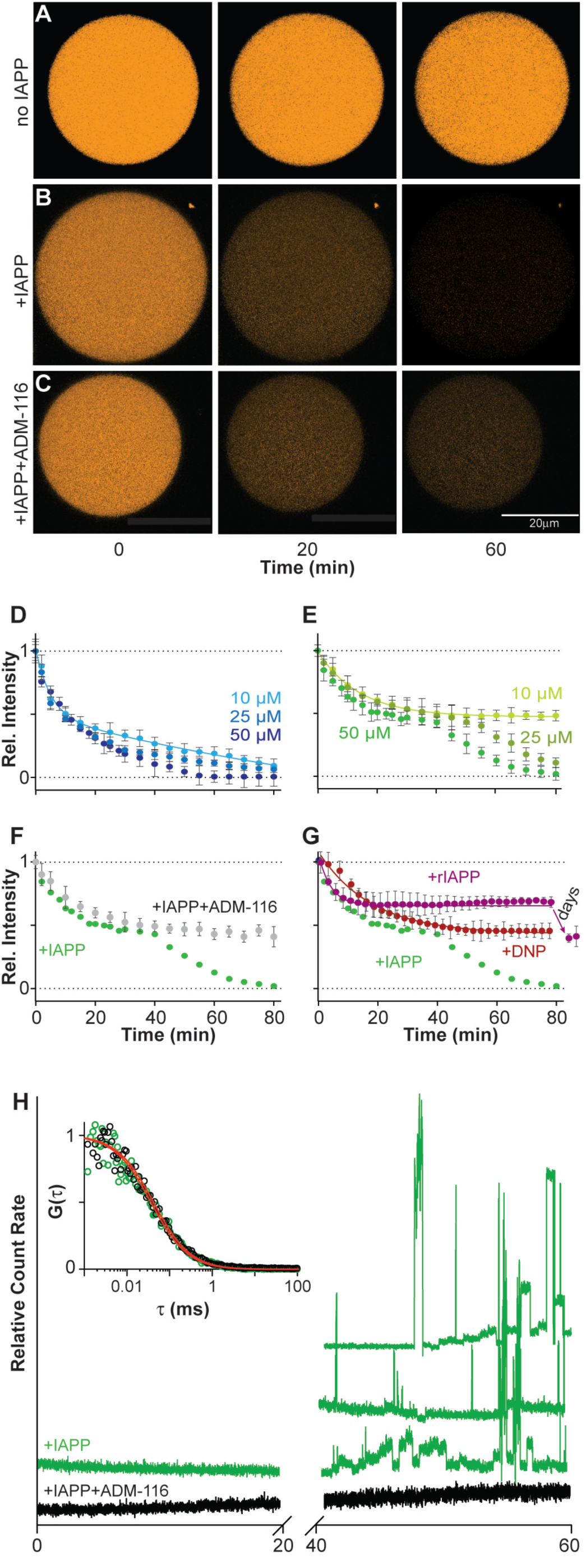
IAPP induces poration of a cell derived membrane. (A-C) Representative confocal imaging of CellTracker Orange labeled GPMVs at the indicated time-points. Shown are single GPMVs observed without added IAPP (A), IAPP added to a ratio of 10:1 (IAPP:lipid) (B), or IAPP and ADM-116 added at a ratio of 10:1:10 (IAPP:lipid:ligand) (C). (D-G) Integration of intensities across repeats of data as in (A-C) used to compute kinetic profiles of leakage from 5 μΜ (phospholipid monomer units) GPMVs. Error bars are standard deviations from three sets of experiments conducted on separate occasions. (D, E) Leakage profiles after addition of the indicated concentrations of IAPP in the presence (D) and absence (E) of 1 mM AB9 (a small molecule fluorescence quencher). (F) Leakage profile at 50 μM IAPP in the presence of equimolar ADM-116. Corresponding data in the absence of ADM-116, (E), is shown as overlay. (G) Leakage profiles upon addition of 10 μM DNP or 200 μM of the sequence variant of IAPP from rats, rIAPP. Corresponding data for 50 μM human IAPP, (E), is shown as overlay. (H) Monitoring the diffusion of fluorescent material escaping the GPMV lumen by FCS. Representative traces of photon bursts are monitored over time in the absence (green) and presence of ADM-116 (black) under conditions matched to (F). Inset: autocorrelation overlaid with fits for data from the first 20 minutes. Right: Three repeats of bursts recorded at time 40 to 60 mins for protein in the absence of ADM-116 are shown. See also Figure S1.

Leakage profiles monitored in the absence of a fluorescence quencher display multiple features. In the absence of quencher, intensity loss by GPMVs in response to IAPP is derived only from the exit of fluorescently labeled cytoplasmic components. Remarkably, two kinetic phases of leakage are observed (Figure 1B and 1E). At 10:1, P:L, the first phase fits to a single exponential decay with a rate constant, 33 ± 2 s^−1^, which is approximately half that observed in the presence of quencher (above). This is a reasonable consequence of removing one (influx of quencher) of the two origins of intensity loss. A much more dramatic difference is the ending of this phase at a plateau in which only 48 ± *4%* of the starting intensity is lost. A second kinetic decay begins after a lag period of 1400 ± 200 s. The profile of this latter phase also fits to a single exponential decay with a rate constant 49 ± 2 s^-1^ and a plateau at baseline. At 5:1, the kinetic constants for leakage are slightly diminished while the lag-phase is more noticeably affected (Figure 1E). At the lowest stoichiometry assayed, 2:1, only the first leakage phase is evident on the timescale of observation (Figure 1E). Plainly, pores formed early are almost independent of IAPP concentration while pores formed later display characteristics that are more dependent upon protein concentration.

The two classes of IAPP pore have distinct sieve size. In the above assays, changes in intensity include contributions from the loss of labeled molecules from the GPMV lumen to the surrounding buffer. To our knowledge, this is the only permeation assay that by design, incorporates heterogeneously sized solutes (all the sulfhydryl containing entities in the cytoplasm). Fluorescence correlation spectroscopy (FCS) was therefore used to assess the diffusion of these fluorescent molecules. During the first of the two kinetic phases of leakage, no large particles are observed (Figure 1H). Correlation analysis is possible and fits are readily made to an analytical form that includes only a single diffusing species. This gives a diffusion constant of 390 ± 25 μm^2^/s. Using the Stokes-Einstein approximation, this corresponds to an average molecular weight of ~700 Da. This may reflect unconjugated dye, or dye conjugated to thiol containing small molecules. In marked contrast, data collection during the second phase of leakage is characterized by large irregular bursts (Figure 1H). This reflects the presence of much larger and heterogeneous species (Elbaum-Garfinkle et al., 2010). Experimental limitations prevent rigorous characterization but particles larger than ~200 kDa is a reasonable estimate. Clearly, the two phases of leakage are characterized by a transition from one in which only small consistently sized holes are present to one in which large heterogeneous defects are formed.

The assemblies of IAPP that give rise to alternative leakage profiles are structurally distinct. Recently, we reported the development of a small molecule, ADM-116, which binds and stabilizes a-helical, membrane bound conformations of IAPP (Kumar et al., 2016a; Kumar et al., 2016b). Moreover, as ADM-116 can cross the plasma membrane and bind to intracellular IAPP, it is uniquely suited for making measurements on GPMVs that can be later matched to assessments inside live cells (see below). Leakage profiles were collected on GPMVs at a P:L of 10:1 and with the addition of ADM-116 equimolar with IAPP (Figure 1C and 1F). The resultant profiles fit to single exponential decay with a rate constant of 34 ± 1 s^-1^ and an amplitude of 52 ± 3 %, comparable to the first phase leakage observed in Figure 1E. FCS measurements of the media show no evidence of release of large heterogeneous species (Figure 1H). Correlation analysis of leaked materials under these conditions yields a diffusion constant of 480 ± 50 μm^2^/s, closely similar to that of materials leaked in the absence of ADM-116 during the first phase. Most significantly, ADM-116 has no effect on the first, small hole leakage process and appears to wholly inhibit the second, large hole leakage process, suggesting that structurally distinct IAPP assemblies are formed on the membrane.

The presence of two classes of pore is a property of IAPP, and not intrinsic to the GPMV based assay. First we consider DNP, a compound known to reduce mitochondrial membrane potential through poration. This compound was recently shown to rescue diabetic animal models (Perry et al., 2015). A concentration of 10 μM DNP induces GPMVs to leak at a rate of 58 ± 8 s^-1^ (Figure 1G). This loss of intensity plateaus with ~50% of fluorescence remaining. By FCS, no large particles emerge from the GPMVs upon leakage (Figure S1). Correlation analysis is readily accommodated by fits to single species giving a diffusion constant, 420 ± 24 μm^2^/s, within error of that observed using IAPP to induce leakage. Importantly, inducing leakage using DNP does not reveal a two-phase process and indeed more closely resembles leakage induced by IAPP when in complex with ADM-116. Second, the sequence variant of IAPP from rat is routinely used as a control peptide as it is not amyloidogenic and requires highly elevated concentrations to display even weak cytotoxicity (Magzoub and Miranker, 2012). At 200 μM (P:L=40:1), rat IAPP induced leakage is observable on the same timescale as human IAPP (Figure 1G). Some additional leakage is apparent, plateauing after a day, but total loss of intensity is ~50%, with no further changes detected even after 4 days (Figure 1G). The contrasting behavior of human IAPP with DNP and with rat IAPP clearly shows that the two phases of human IAPP induced leakage are specific to this variant of the peptide.

### Intracellular loss of membrane integrity

IAPP localization to mitochondria is correlated with depolarization. IAPP induced toxicity in cells has been associated with several cellular membranes, most notably mitochondria. This is observed under conditions where IAPP is expressed (Gurlo et al., 2010) or introduced exogenously into the culture medium (Kumar et al., 2016b; Magzoub and Miranker, 2012). Mitochondrial uncoupling in response to IAPP has been previously reported (Soty M, 2011), but without assessment of correlation with IAPP localization. Here, we have used the membrane potential sensitive dye, JC-1, to monitor mitochondrial polarization in live INS-1 cells in the presence IAPP doped with 200 nM IAPP labeled with Atto647 (Figure 2A). Addition of 13 μM IAPP to the culture media results in greater than 50% cell death after 48 h (Figure S2). Imaging performed at 24 h shows widespread mitochondrial localization accompanied by loss of polarization (Figure 2B and 2E). Overlap analysis across 50 cells shows ~60% of observable IAPP to be associated with depolarized mitochondria (Figure 2F). By 48 h the extent of loss in membrane polarization is quadrupled (Figure 2B and 2E). At 5 μM IAPP, INS-1 cells experience a reduced cytotoxicity of ~15% at 48 h (Figure S2). Mitochondrial depolarization at 24 h and at 48 h is still evident (Figure 2C and 2E) but greatly diminished relative to the more toxic condition. IAPP colocalization to depolarized mitochondria is still evident albeit reduced to ~20% of the total IAPP (Figure 2F). In contrast, IAPP overlap with ER is 5% and 7% for these two conditions respectively (Figure 2H, 2I and 2J). Thus, localization of IAPP to mitochondria appears to be strongly associated with depolarization. Moreover, it is the extent of depolarization that correlates most strongly with the toxic condition.

**Figure 2:**
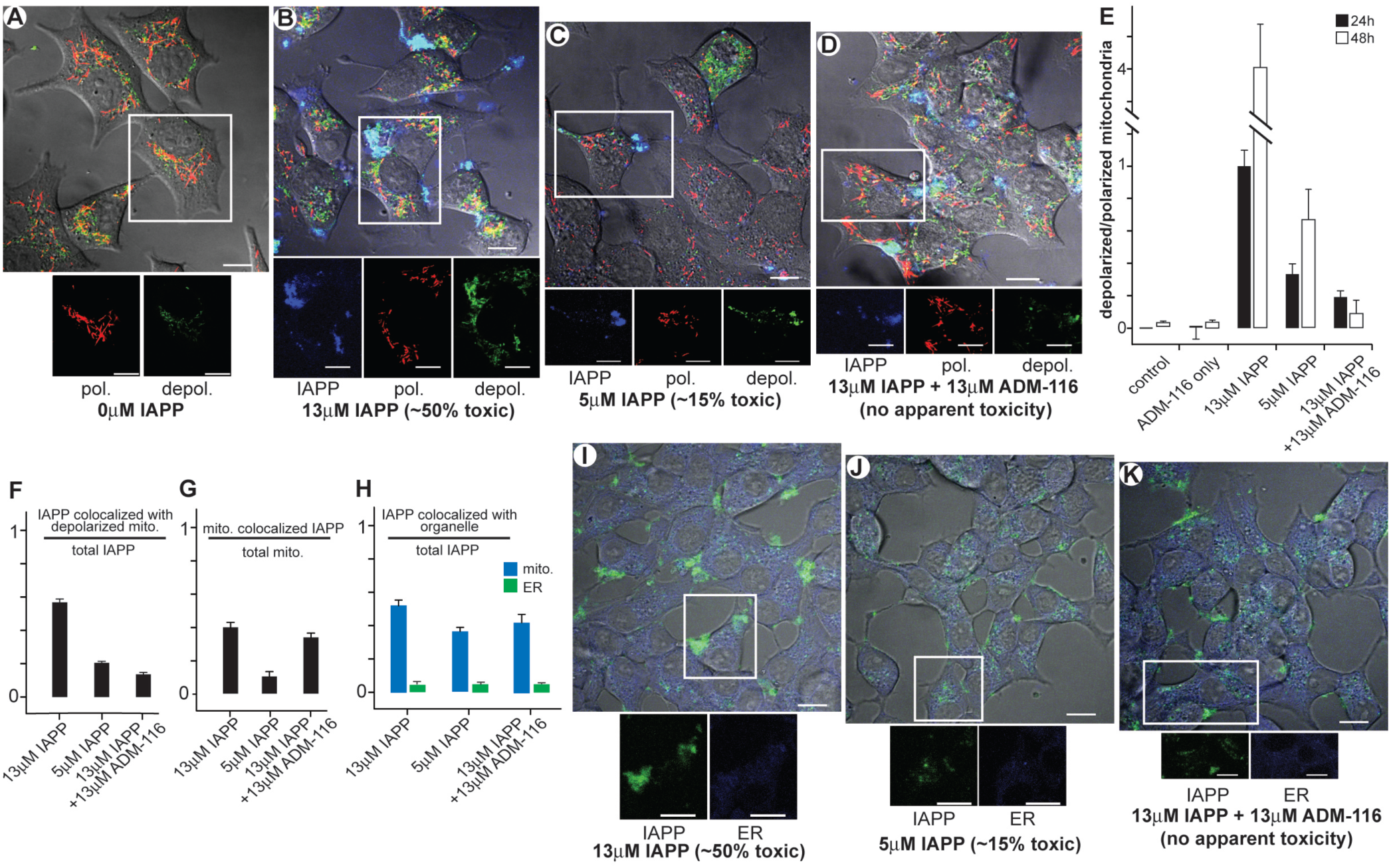
Mitochondria colocalized IAPP is associated with depolarization. (A) Representative INS-1 cells stained with JC-1 marker. Two fluorescence channels in red and green correspond to polarized and depolarized mitochondria respectively. These are shown merged with the differential interference contrast (DIC) image. A region of interest (ROI) is chosen to show channels individually. (B) Representative cells exposed to toxic (13 μΜ) concentrations of IAPP doped with 200 nM IAPP_A647_ for imaging. Cells were stained with JC-1 marker 24 h after IAPP. The three channels, IAPP_A647_ (blue), polarized (red) and depolarized (green) mitochrondria are shown merged with the DIC image. A region of interest (ROI) is chosen to show channels individually. (C) As in Figure 2B but at 5 μΜ IAPP. (D) As in Figure 2B, but with the addition of 13 μΜ ADM-116 three hours after addition of IAPP (to ensure IAPP is intracellular). (E) Image overlap statistics for Figure 2A-D showing ratio of depolarized/polarized mitochondria at the indicated conditions and incubation time. (F) Image overlap statistics for Figure 2B-D showing fraction of total IAPP overlapping depolarized mitochondria. (G) Image overlap statistics for Figure 2B-D showing fraction of total mitochondria colocalized with IAPP. (H) Image overlap statistics for Figure 2B-D and Figure 2I-K, showing fraction of total IAPP overlapping with mitochondria and ER respectively. (E-H) For each measurement, a total of 50 cells from 3 biological replicates were selected for analysis. Error bars are standard deviations from each condition’s set of cells. (I-K) As in Figure 2B-D, but monitored using ER-Tracker instead of mitochondrial polarization marker and doping IAPP with 200 nM IAPP_A488_. Scale bar for all images is 20 μm. See also Figures S2 and S3.

Cytotoxic rescue by ADM-116 significantly diminishes the correlation of IAPP with mitochondrial depolarization. Under our standardized rescue condition, 13 μM ADM-116 added to culture media 3 h after uptake of 13 μM IAPP, depolarization at 24 h is markedly reduced (Figure 2D and 2E) as is the overall apparent extent of IAPP association with depolarized mitochondria (Figure 2F). The capacity of ADM-116 to reduce mitochondrial depolarization remains potent through 48 h incubation (Figure 2E). Importantly, rescue of IAPP induced toxicity with ADM-116 does not significantly change the overall amount of protein localized with mitochondria (Figure 2H), nor the fraction of mitochondria with associated IAPP (Figure 2G). Moreover, neither rescue with ADM-116 nor a reduction in toxicity using less IAPP wholly eliminates depolarization (Figure 2E). For comparison, experiments were performed in which mitochondria were manipulated instead by incubating INS-1 cells with DNP. At routinely used concentrations (5 μM and 10 μM), we find DNP is weakly toxic and JC-1 measured mitochondrial polarization is unaffected (Figure S3). This is analogous to reports using yeast that show that sub-toxic levels of DNP induce changes in mitochondrial respiration without significant loss of membrane polarization (Antonenko et al., 2013). It is therefore plausible that small pore leakage induced by IAPP in the presence of ADM-116 is not sufficient to cause depolarization of cellular mitochondria.

### Coarse grain structural analysis via intermolecular FRET

Structural analysis of IAPP oligomers was performed via intermolecular FRET. Briefly, single labeled IAPPs are prepared in which either a donor (Alexa 488, IAPP_A488_) or acceptor (Atto 647, IAPP_A647_) fluorophore is covalently attached to the N-terminus of IAPP. We deviate from typical intermolecular FRET protocols by performing experiments under conditions where labeled IAPP is mixed with a high proportion of unlabeled IAPP. This approach, which we term “diluted-FRET”, draws observation to the subset of IAPP oligomers that are sufficiently large to harbor at least one donor and acceptor pair under the diluted condition.

GPMV associated IAPP forms a discrete species when in complex with the structure specific small molecule binder, ADM-116. GPMVs were prepared as for leakage studies above, but without labeling the lumen contents. Under conditions matched to those that inhibit the second porated state (P:L:ADM-116=10:1:10) (Figure 1F), FRET is readily apparent at an empirically determined dilution of 1:1:500 (donor:acceptor:unlabeled IAPP) (Figure S5). At this dilution, hundreds of IAPP must be in proximity in order to yield significant observation of FRET. Confocal imaging of GPMVs show formation of localized puncta within 2’. These appear to behave as foci from which amorphous assemblies grow (Figure 3). This is in marked contrast to observations made using synthetic GUVs which show IAPP forming a uniform distribution on the membrane (Gao and Winter, 2015; Kegulian et al., 2015; Seeliger et al., 2012). In addition, ADM-116 neither prohibits nor displaces IAPP from the GPMV surface, consistent with mitochondria localization studies described above. Using a toroidal region of interest (ROI) that is predominantly occupied by membrane, time dependent changes to FRET_eff_ are suggestive of progressive transitions between discrete forms (Figure 3C). Qualitatively over the first ~24’, two apparent species are sampled before the distribution reaches a third at FRET_eff_ = 0.47 ± 0.03. Progression to higher FRET_eff_ continues but stabilizes after t=40’ with FRET_eff_ = 0.62 ± 0.03. Thus, under conditions in which only first phase poration is observed, changes in oligomeric IAPP assemblies occurs on a comparable timescale.

**Figure 3:**
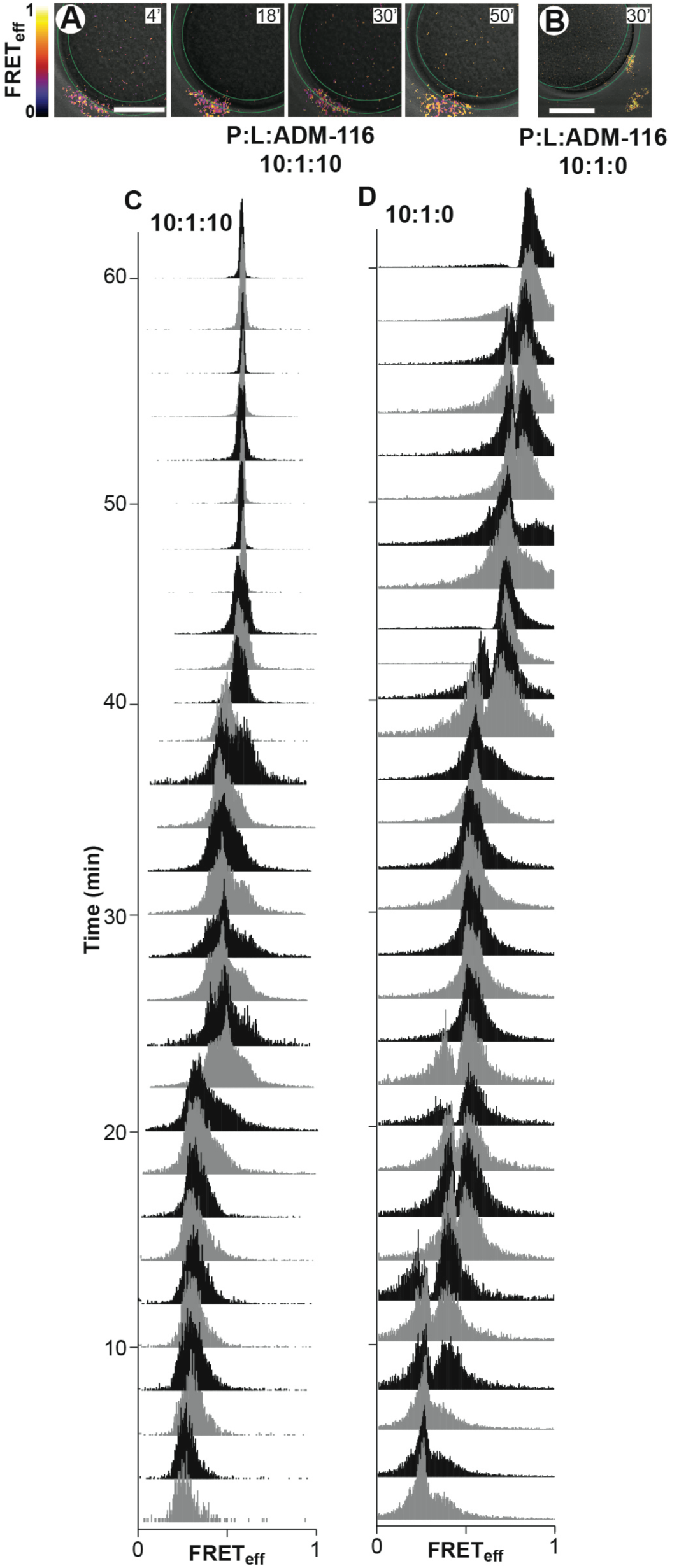
Time resolved conformational changes of IAPP on GPMVs match leakage profiles. Time dependent changes to intermolecular FRET between IAPP_A_488 and IAPP_A_647 (in the presence of unlabeled IAPP, 1:1:500) imaged on GPMVs and shown as a heat map merged with corresponding DIC images. Below these, image statistics are shown as histograms of pixel counts. (A, B) Representative time points of a single ring shaped ROI (green) that predominantly collects contributions from the membrane. GPMVs at 5 μΜ (phospholipid monomer units) incubated with IAPP, lipid and ADM-116 at the indicated ratios. Scale bar, 10 μm. (C, D) FRET_eff_ histograms from single optical sections of a single GMPV monitored over time after exposure to IAPP and ADM-116 at ratios of 10:1:10 (C) and 10:1:0 (D) protein:lipid:ADM-116. Images were recorded every 2’ using ring-shaped ROIs that encompass the membrane. See also Figure S4.

Time resolved analysis of diluted-FRET of IAPP, under conditions resulting in complete leakage of GPMVs, indicates the presence of six species (Figure 3D). At protein to lipid ratios of 10:1:0 (no small molecule), the first oligomers appear just as they did in the presence of small molecule, i.e. as puncta within the ~2’ deadtime of measurement. Over the timescale associated with observation of leakage, five additional assemblies emerge, each with a discrete, approximately Gaussian distribution of FRET_eff_. The expected behavior for growth resulting from monomer or small oligomer additions to these assemblies would be horizontal shifting of the FRET_eff_ peak position. This is not observed. Instead, no species appears without another disappearing. This occurs in succession, with each peak showing greater FRET_eff_ than its predecessor. Moreover, overall counts increase with increasing FRET_eff_. These two observations suggest that the progression in FRET_eff_ is a result of stepwise merging of smaller oligomers. Comparable changes are observed, albeit with different average FRET values, for an alternate donor/acceptor pair (Figure S4A). The invariance with choice of fluorophores and the coincidence of the timescale with that observed for leakage (where all IAPP is unlabeled) are independently consistent with the fluorophore not contributing significantly to the structures sampled by IAPP oligomers. Plainly, the conformational landscape of membrane associated IAPP contains multiple transitions that occur on timescales associated with membrane integrity loss.

The two classes of pores formed by IAPP on GPMVs correspond to distinct species of oligomer distinguishable by their diluted-FRET_eff_. Combining repeats of the above data across multiple GPMVs results in robust characterization of individual species (Figure 4). Under our standard condition for observing two state leakage (P:L:ADM-116=10:1:0), summation over the first 34’ reveal four diluted-FRET distribution peaks with FRET_eff_ at 0.21 ± 0.07, 0.36 ± 0.04, 0.41 ± 0.08 and 0.58 ± 0.02 respectively (Figure 4A*i*, 4A*ii* and 4A*iii*). We observe the first three FRET_eff_ peaks to dominate over a period of time in which only small molecule leakage is observed (Figure 1E, H). Analysis between 36’ and 46’, i.e. the time period over which large molecule leakage initiates, continues to be dominated by ensembles that were previously present (Figure 4A*iv*), although with a shift to FRET_eff_ = 0.58 as the dominant peak. Summation across 48’ to 60’ show a marked shift to two high FRET_eff_ ensembles at 0.78 ± 0.04 and 0.87 ± 0.01 (Figure 4A*v*) with the former dominant over the majority of this leakage phase (Figure 3F). Detection of six structural forms over a timescale in which only two leakage processes are evident suggests that not all oligomer growth effects change to poration relevant membrane structures (see discussion). Under conditions in which the second porous phase is eliminated or delayed, i.e. in the presence of ADM-116 (Figure 4B) or at reduced concentrations of protein (P:L=5:1:0) (Figure 4C) respectively, the final FRET_eff_ distributions are each dominated by a peak whose position corresponds to that of the fourth peak observed at 10:1:0 (P:L:ADM-116). In other words, conditions that affect large but not small hole poration (Figure 1), exclusively affect access to the last two of the six species that are apparent using diluted-FRET.

**Figure 4:**
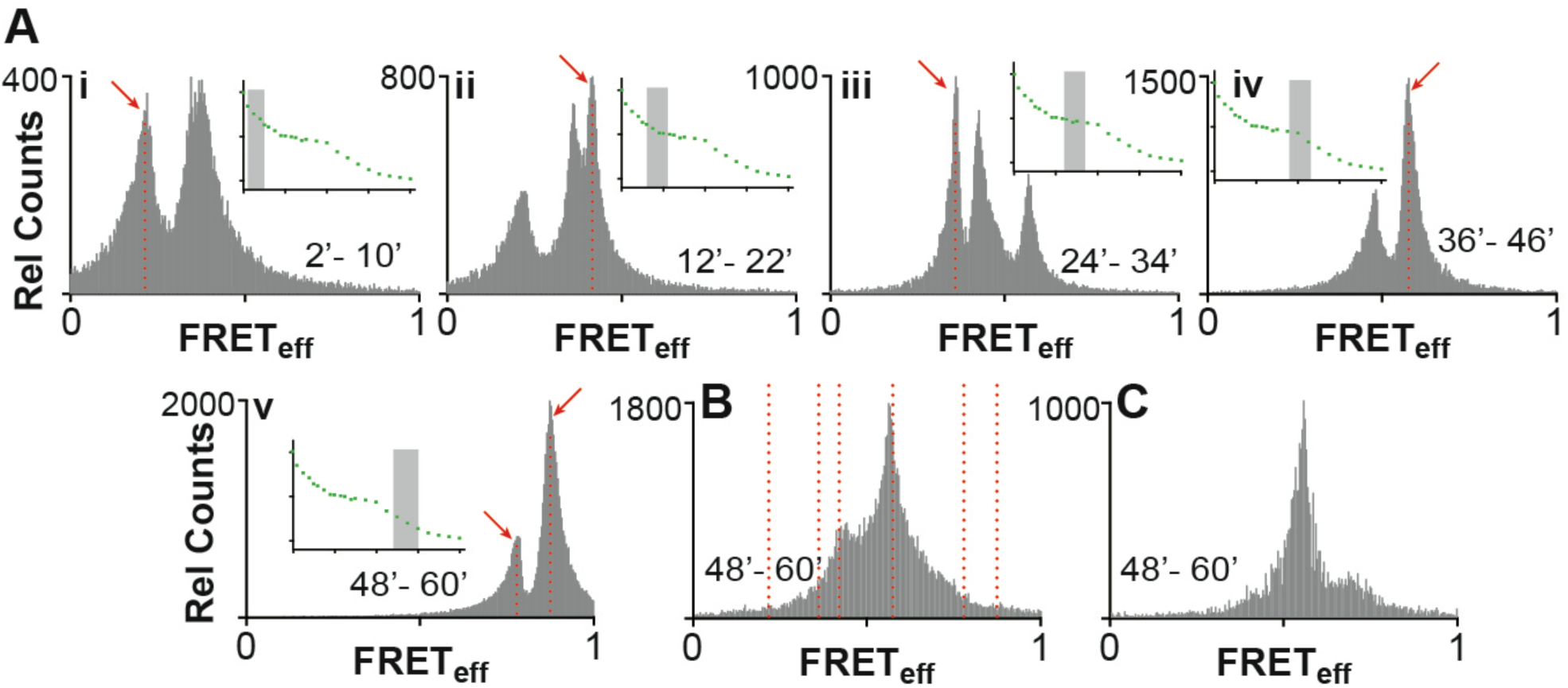
Global changes to IAPP oligomers on GPMVs. Intermolecular FRET histograms summed over a minimum of three GPMVs, five optical sections for each GPMV and the indicated periods of time. Conditions are matched to those shown in Figure 3. (A) Diluted-FRET at 10:1:0 IAPP:lipid:ADM-116 (Histograms are shown *(i-v)* corresponding to summation across the indicated time period. Red arrows indicate peaks that define FRET_eff_ values for species referred to in the main text and the locations of dotted red lines in (B) and Figure 5A. Inset: Schematic of leakage profile collected under matched conditions (Figure 1F). Grey rectangles show the summed time period indicated on the associated FRET_eff_ histogram. (B, C) As in (A), but corresponding to IAPP, lipid and ADM-116 at ratios of 10:1:10 (B) and 5:1:0 (C) respectively. See also Figures S5.

The two highest FRET_eff_ species observed on GPMVs do not correspond to canonical IAPP fibril structures. In the above analysis (Figure 3A), structures are first evident at ~2’ and enlarge throughout the observation time regardless of the presence or absence of ligand. In the absence of ADM-116, determinations of FRET_eff_ using ROIs that encompass IAPP assemblies near but not on a GPMV surface (Figure 5A) give a distribution wholly consistent with surface associated IAPP (Figure 4Av). In marked contrast, the FRET_eff_ distribution of fiber controls are broad and possess a single peak with a FRET_eff_ that is greater than what is observed on GPMVs (Figure 5A). The distribution is Poisson-like with significant FRET_eff_ counts throughout the range of zero to one. This clearly distinguishes the fiber control from leakage competent oligomers. In addition, IAPP oligomers on and near GPMVs do not enhance the fluorescence of the amyloid indicator dye, Thioflavin T (LeVine, 1993) (Figure 5B). Systematic error can arise as GPMVs independently contribute to the fluorescence intensity of ThT, and intensity changes in ThT can also result from changes in aggregate size. To accommodate this, an internal control, IAPP_A647_, was included so that ThT intensity could be renormalized on a pixel by pixel basis. The renormalized response of GPMV generated IAPP aggregates are 6-fold lower than that observed for the fiber control (Figure 5D). Therefore, the oligomer structures of IAPP associated with membrane leakage cannot be predominantly formed from canonical amyloid fibers.

**Figure 5:**
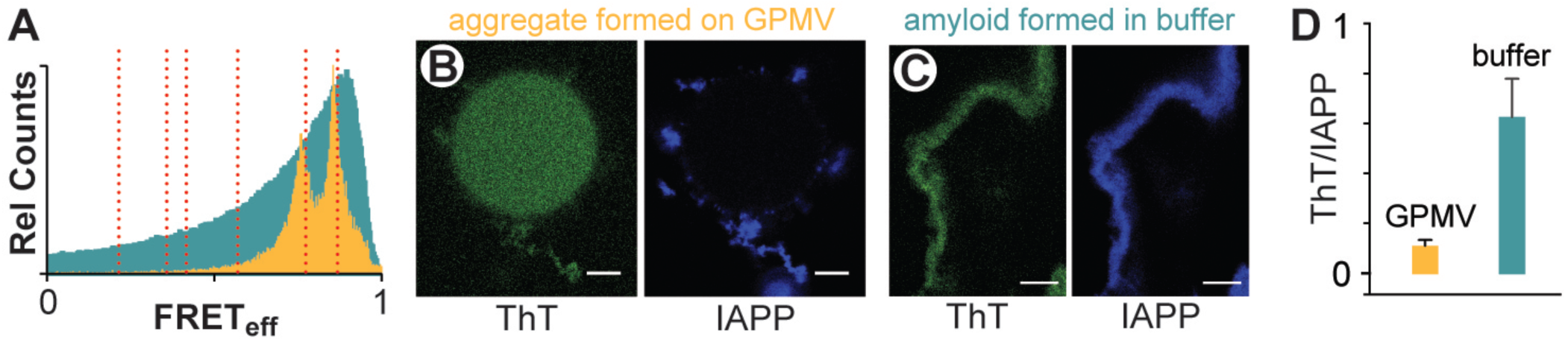
IAPP oligomers on GPMVs do not have a canonical amyloid fiber structure. (A) Intermolecular FRET_eff_ statistics from 50 ROIs of aggregates prepared using IAPP_A488_, IAPP_A647_ and unlabeled IAPP (1:1:500). GPMV associated aggregates (orange) were prepared under conditions of 10:1:0 IAPP:lipid:ADM-116 (matched to Figure 4A) with ROIs taken at 60’ timepoints and at locations distal from the GPMV surface. Data from amyloid fibers prepared as control in aqueous buffer are shown in cyan. (B, C) Confocal imaging of IAPP aggregates prepared from IAPP doped with IAPP_A647_ (5 00:1) and further stained with the amyloid indicator dye, ThT. Two channel image pairs are shown for ThT (green) and IAPP_A647_ (blue) respectively. Aggregates were prepared either in the presence of GPMVs (B) or in buffer (C). (D) Imaging statistics for data as in (B, C). Presented as the ratio of ThT intensities divided by IAPP_A647_ intensities using ROIs away from the GPMV surface. Error bars are standard deviations from three sets of experiments conducted on independent occasions.

Diluted FRET reveals large-*N* oligomers inside cells that uniformly adopt a single overall structure when toxicity is rescued by ADM-116. To parallel *in vitro* measurements on GPMVs, INS-1 cells were incubated with 10 μM IAPP, corresponding to 50% cell toxicity (Figure S2). As previously established for cellular rescue experiments, 10 μΜ ADM-116 was added to culture media 3 h after IAPP to ensure the protein was intracellular prior to addition of small molecule (Kumar et al., 2016b). Imaging conducted at 24 h (Figure 6A) shows intracellular, intermolecular FRET to be widespread indicating that large *N* oligomers are abundant. The ideal, empirically determined doping ratio for these cellular experiments is 1:1:100 (donor:acceptor:unlabeled IAPP), in contrast to FRET measurements on GPMVs performed at 1:1:500, suggesting that the intracellular oligomers are smaller than those observed on GPMVs. The FRET_eff_ histograms of individual cells are directly comparable to averages conducted over many cells (Figure 6B) and show a single, narrow and time invariant distribution with a peak efficiency of 0.60 ± 0.07. After 48 h, this distribution has moved only slightly. Analysis using a different FRET pair, IAPP_A488_ and IAPP_A594_, give similar results (Figure S4B). This suggests the fluorophores do not contribute significantly to the structures giving rise to the observation. Plainly, a single, long-lived discrete assembly of intracellular IAPP is formed upon cytotoxic rescue by ADM-116; a result comparable with that observed on GPMVs (Figure 3C and 4B).

**Figure 6:**
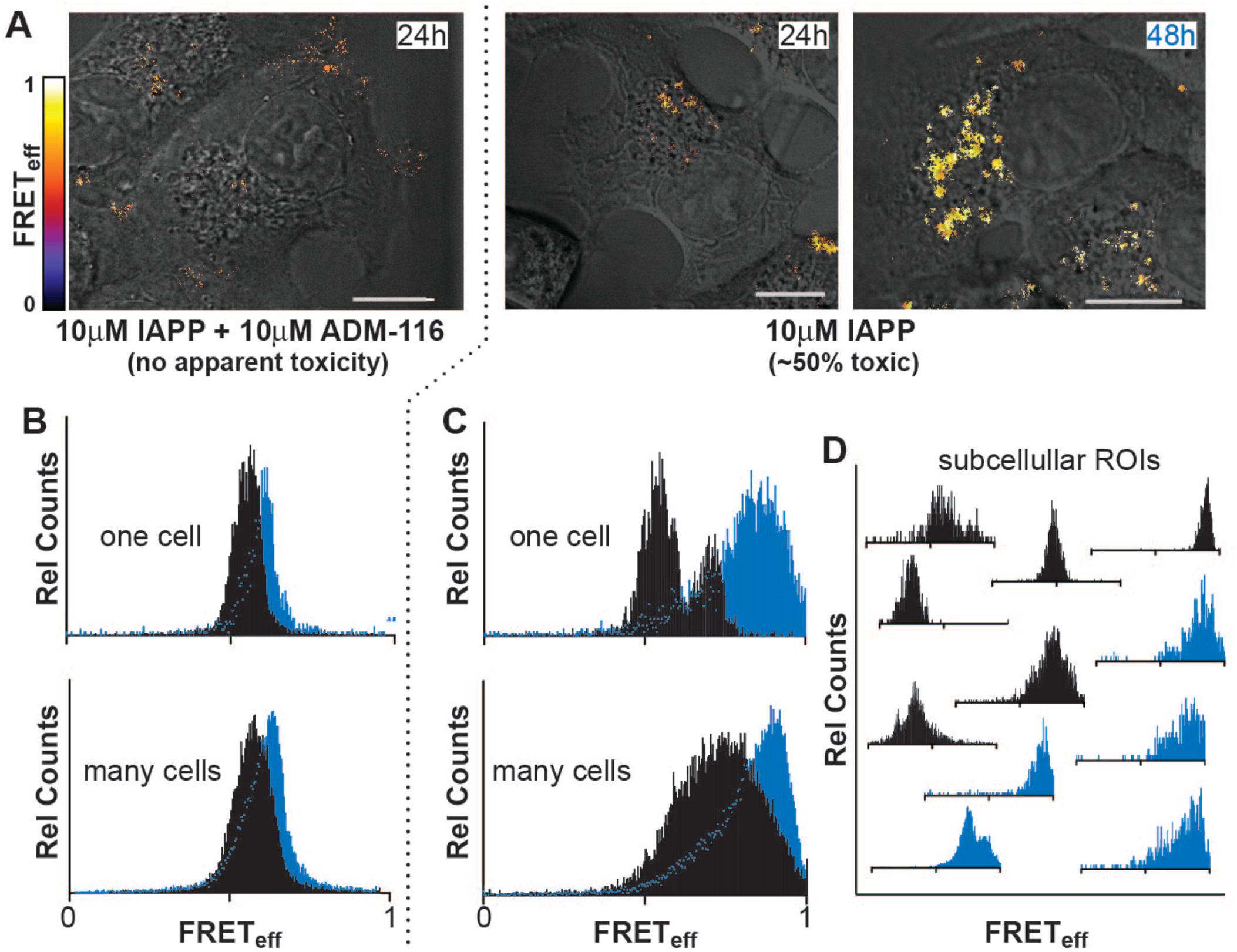
IAPP adopts a specific conformation in cells upon rescue. (A) Intermolecular FRET between IAPP_A488_ and IAPP_A647_ (in the presence of unlabeled IAPP, 1:1:100) imaged in live INS-1 cells under the conditions indicated. Pixel-based FRET analysis of representative ROIs are shown as a heat map merged with corresponding DIC images. Scale bar, 10 μm. (B-D) Histograms of pixel counts for FRET_eff_ images as in (A). (B) FRET_eff_ histograms for cells at 24 h (black) and 48 h (cyan) for the ADM-116 rescued condition (A). Analysis for a representative single cell is shown above the sum of data collected from 50 cells from 3 independently conducted experiments on each of 3 different days. (C) Analysis as in (B), under conditions (10 μΜ IAPP) in which ~50% toxicity is measured colorimetrically (Figure S2). (D) Analysis as in (C), but for representative ROIs within single cells. See also Figures S2 and S4.

Intracellular oligomers of IAPP evolve to high-FRET_eff_ states under conditions that are cytotoxic. Diluted-FRET was measured using cytotoxic levels of IAPP (10 μM) at 24 h and 48 h without rescue using ADM-116. Cumulative analysis of cell images taken at 24 h reveal a broad range of FRET_eff_ dominated by values much greater than 0.5 (Figure 6C). Individual cells (Figure 6C) as well as small sub-cellular ROIs (Figure 6D) show recurring and relatively narrow distributions at a small number of reproducible values of FRETeff. This is inline with observation of discrete species observed on GPMVs (Figure 4). The breadth of the cumulative distribution (Figure 6C) may therefore be a consequence of summing contributions from discrete rather than a continuum of species. After 48 h, cells show significant morphological signs of dysfunction. This is accompanied by FRET distributions shifting to markedly higher efficiencies (Figure 6C and 6D). Plainly, IAPP oligomer structures inside cells mimic those sampled on GPMVs. Specifically, given cytotoxic conditions on cells, or leakage conditions on GPMVs, a time dependent evolution of IAPP oligomers to high FRET_eff_ is apparent. When an IAPP specific ligand is used to rescue cells or inhibit leakage, an intermediate state is stabilized.

## Discussion

The task of ascribing structure to function is unusually challenging for pre-amyloid oligomeric toxins. The structures are plastic, oligomeric size is heterogeneous and often there is a dynamic exchange between aqueous and membrane-associated forms. Here, these challenges have been overcome through use of a structure-specific small molecule ligand and comparative measurements in cell-derived model membranes and live-cell imaging.

The use of GPMVs enabled the identification of two forms of membrane permeation. This is apparent from two kinetic phases in the leakage profiles (Figure 1E) each with a characteristic size distribution for the escaping molecules (Figure 1H). Small molecule binding to IAPP inhibits the second, but not the first phase of leakage (Figure 1F). A parallel can be drawn between these results and the mitochondrial depolarization measurements in cells (Figure 2), where the small molecule ligand reverses the depolarization seen in the presence of IAPP, without displacing IAPP from mitochondria (Figure 2G). This observation is significant, as it shows localization of IAPP to mitochondria to be necessary but not sufficient for toxicity. Instead, it appears that the structure of IAPP at the mitochondria is a more relevant indicator of toxic effect. This is supported by the observation that on GPMVs and on mitochondria, halting of leakage/depolarization and toxic processes are each correlated with stabilization of an intermediate oligomer (Figure 3C and 6B).

These results serve as a basis for us to propose that nucleated conversion of small pore permeable states to large pore permeation is the origin of gains in toxic function by IAPP. This is an important distinction as poration is not necessarily toxic. Indeed, recent work with DNP shows that poration of mitochondria can be cytoprotective (Perry et al., 2015). Using GPMVs, we find that DNP mediated poration (Figure 1D) results in single kinetic phase that does not go to baseline, consistent with small hole poration induced using IAPP (Figure 1E, 1H and S2). While these results clearly demonstrate a link between large hole poration of mitochondria and cell toxicity, they leave open the exact mechanism of toxicity. Toxicity may occur directly at the mitochondria, or, alternatively, it may be one of several possible consequences, such as oxidative stress, that can result from mitochondrial poration (Costes et al., 2013a).

The diluted-FRET measurements can be leveraged to delineate some of the structural features that distinguish toxic from non-toxic poration. For example, the IAPP oligomers responsible for membrane disruption consist of scores to hundreds of peptides. As the overwhelming majority of IAPP (1:1:100 and 1:1:500 donor:acceptor:unlabeled) present in these measurements is unlabeled, it is highly improbable that small-*N* sized structures would give rise to FRET. Consider an extreme in which only dimers are formed. A binomial distribution of labeled IAPP among these dimers would result in donor/acceptor pairs occurring < 0.1% of the time. Instead, FRET is abundant in our images, with at least 70% of acceptors contributing to our observations (see Methods). At 1:1:100 (donor:acceptor:unlabeled protein), oligomers with *N* > 30 are required to observe this degree of prevalence of FRET. At 1:1:500, our estimate rises to *N* > 150.

The membrane-disrupting states of IAPP appear to not only be large, but also dynamic. This is suggested by the diluted-FRET measurements in both live cells and on GPMVs. Notably, the measured FRET distributions are very narrow. A narrow distribution would be expected for well-defined states where the donor and acceptor fluorophores are held at a constant distance apart. The combinatorial distribution of labeled IAPPs within a large-*N* oligomer prohibits this possibility. Narrow FRET_eff_ distributions therefore must be the result of signal averaging. Averaging may arise from the presence of many static oligomers with different FRET efficiencies present within a confocal section. Alternatively, averaging might be a consequence of rapid rearrangement of IAPP within oligomers. IAPP amyloid fibers (Figure 5A) allow for delineation between these two possibilities. Fibers are not internally dynamic and are highly dense assemblies of IAPP. In striking contrast to the narrow FRET efficiency peaks measured in cells and an GMPVs, the FRET measured in fibers display a broad, asymmetric FRET_eff_ peak with significant counts occurring across the full span of 0 to 1 (Figure 5A). This observation supports a model where time-averaging of the FRET signal is occurring for membrane associated IAPP. Dynamic rearrangement within oligomers is the most plausible origin for this observation.

The transition between small and large pore states reflects a change in oligomer topology, and/or the conformation of IAPP within the oligomer. Consider that a reasonable volume for one, wholly compacted IAPP is ~4000 Å^3^. At this density and at a donor:acceptor:unlabeled ratio of 1:1:500 (as used for GPMV experiments), a uniformly random distribution of *N*-monomers within a sphere yields a FRET_eff_ that can not exceed ~0.27 (Figure 7). This is clearly smaller than all but one of the observed peaks (Figure 4A). Increasing simulated protein density would allow higher FRET_eff_, but would not be physical. Furthermore, specifically stabilizing a membrane bound α-helical form of IAPP using a small molecule differentially affects alternative phases of poration (Figure 1). These observations suggest oligomeric IAPP possess sufficiently well-defined structure and that changes in structure can affect function.

**Figure 7:**
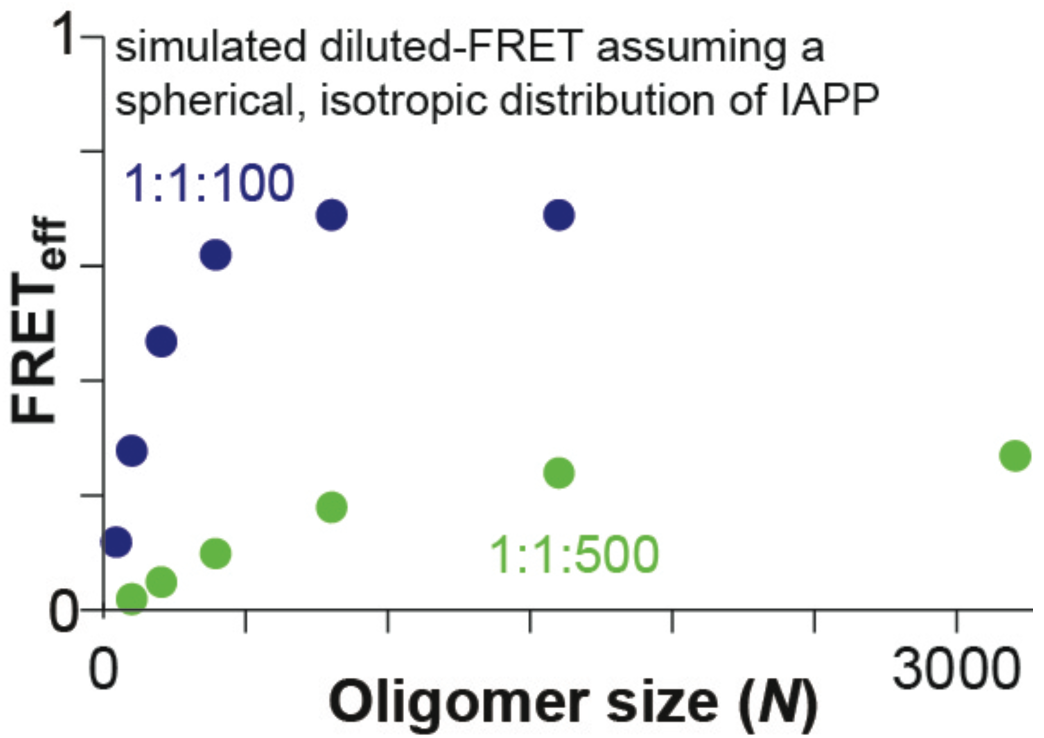
A model for oligomer growth. Simulated average FRET_eff_ for a dynamic, isotropic spherical distribution of an IAPP oligomer of size *N.* Each IAPP molecule is assumed to occupy 4000 Å^3^, a Förster distance of 52 Å, the indicated ratio of (donor labeled IAPP):(acceptor labeled IAPP):(unlabeled IAPP), and the indicated oligomer size.

The evolution of the amyloid-hypothesis into the oligomer-hypothesis was pioneered, in part, by the immunostained light-microscopy observation of membrane associated puncta (Gurlo et al., 2010). The current work goes further by placing limits on the size of assemblies relevant to the development of IAPP-mediated toxicity. A recent effort with most bearing on our work is a study in which SDS was used as a membrane model (Bram et al., 2014). The resultant IAPP oligomer was toxic, chromatographically stable, discrete in size (~90kDa), rich in α-helical secondary structure, shown to enter cells and localize to mitochondria. Remarkably, anti-sera from diabetic patients but not healthy controls were cross reactive with this oligomer. The 90kDa size and subcellular localization observed in that construct is consistent with the observations reported here. Moreover, we have previously shown that ADM-116 stabilizes an α-helical membrane bound form of IAPP (Kumar et al., 2016b). Our current study does not exclude the possibility that large species observed on membranes requires initial steps of dimer through hexamer formation as suggested by other groups including our own (Young et al., 2014) (Abedini et al., 2016)(Nath et al., 2011). However, our study does make clear that the oligomers associated with sub-cellular membranes responsible for gains-in-toxic function are much larger and dynamic.

This dynamic picture of oligomers prompts a reconsideration of the role small molecule interactions can play in protein assemblies involving unstructured or partially structured proteins. Our work here has shown not only the existence of multiple membrane-associated oligomeric states, but that, remarkably, these states are sufficiently well-defined as to be distinguishable by structure-specific small-molecule binding. In amyloid disease systems, membrane poration has been reported for Aβ from Alzheimer's, α-Synuclein from Parkinson’s disease, and PrP from spongiform encephalopathy – each of these proteins contain disordered regions. In functional systems, dynamic assemblies of membrane proteins have been observed for Nephrin (Banjade and Rosen, 2014) and more recently LAT (Su et al., 2016), which forms micron sized clusters during T-cell activation. The small molecule ADM-116 is therefore proof-of-concept to the idea that changes to dynamic, partially ordered protein oligomers are addressable by small molecule development for basic and translational research.

Numerous mechanisms of poration by IAPP and other amyloidogenic proteins have been proposed (Last et al., 2013). These range from structured proteins forming well defined holes (Laganowsky et al., 2012), to protein mediated detergent-like abstraction of phospholipid (Jayasinghe and Langen, 2007). Given the large number of IAPP molecules required for poration, however, the former is unlikely, while apparent GPMV robustness during leakage renders the latter unlikely. A speculative description more consistent with our data is that poration by IAPP is a consequence of a loose association of peptide and lipid that forms a discrete and semi-permeable subdomain within the bilayer. This is conceptually similar to what has been reported for anti-bacterial toxins (Gregory et al., 2008) and referred to as a chaotic-pore (Axelsen, 2008). Such a model allows for pore size to not be affected by the overall size of the membrane embedded oligomer. Moreover, direct protein-protein interactions would have a diminished role as phospholipids are an integral part of the phase. With this model, the conversion of small poration to large poration is simply a phase transition. Indeed, phase-separation was already a property established by us for IAPP in solution (Padrick and Miranker, 2002); (Rhoades and Gafni, 2003). We are only now coming to appreciate that this property is meaningful in a cellular membrane context. The work here goes further by showing phase separation to be manipulable by a protein specific ligand. This validates the idea that small molecules can be used to specifically manipulate other partially ordered systems.

## Author contributions

M.B., E.R, and A.D.M. designed the project and experiments. M.B. performed the experiments. M.B., E.R., and A.D.M, analyzed the data and wrote the manuscript. S.K. synthesized the oligoquinoline amides.

## Acknowledgements

This research was supported by NIH GM094693 (to A.D.M.), GM102815 (to A.D.M. and E.R.) and an American Diabetes Association mentor-based postdoctoral fellowship to M.B. We thank Prof. M. Rosen for critical reading of this work. We thank Alex Miranker for assistance with programming diluted-FRET simulations and Dr J. Wolenski for technical assistance with microscopy.

## METHODS

### Materials

Chemicals were purchased from Sigma Aldrich (St. Louis, MO) unless otherwise specified. Thioflavin T (ThT) was purchased from Acros Organics (Fair Lawn, NJ) and islet amyloid polypeptide (IAPP) from Elim Biopharmaceuticals (Hayward, CA, USA). ADM-116 was previously prepared as described (Kumar et al., 2016a). IAPP stocks were prepared by solubilizing ~2mg protein in 7 M guanidinium hydrochloride. The solution was filtered (0.2 micron) and transferred to C-18 spin column, washed twice with water followed by a wash of 10% acetonitrile, 0.1% formic acid (v/v) and then eluted into 200 μL of 50% acetonitrile, 0.1% formic acid (v/v). The concentration of IAPP was determined by OK at 280 nm ( ε = 1400 M^-1^cm^-1^). The solution was then divided into single use aliquots (20-50 μL), lyophilised, and stored at −80°C. Stock solutions of IAPP were prepared with water from these aliquots. Alexa 488 carboxylic acid succinimidyl ester (A488), Atto 647N succinimidyl ester (A647) and Alexa 594 succinimidyl ester (A594) dyes were purchased from Life Technologies (Carlsbad, CA). IAPP labelling was prepared as described previously (Magzoub and Miranker, 2012). Briefly, IAPP was incubated with dye, succinimidyl ester on a MacroSpin column for 4 h. Labelled IAPP was eluted from the MacroSpin column with 50% acetonitrile/0.2% formic acid solution. This was then diluted with 7 M guanidinium hydrochloride solution to a total organic content of <5%. Labelled protein was then purified by reverse-phase high-performance liquid chromatography, and identity was confirmed by mass spectrometry. Aliquots were lyophilized and stored at –80°C. Stocks at 100 *μ*M in water were prepared and immediately before use. IAPP control fibers were prepared by incubating 50 *μ*M hIAPP and 100 nM IAPP_A647_ in 50 mM sodium phosphate buffer, 100 mM KCl, pH 7.4, for ~24 h. Fibers were pelleted at 21,000 *g* for 30 min and resuspended three times using water..

### Giant Plasma Membrane Vesicles

GPMVs were isolated from INS-1 cells as previously described (Schlamadinger and Miranker, 2014). Briefly, cells were plated in 35 mm dishes and cultured for 48 h, washed with a 10 mM HEPES, 150 mM NaCl, 2 mM CaCl_2_ (pH = 7.4) twice and then exposed to 10 mM N-ethyl maleimide (NEM, Sigma Aldrich, St. Louis, MI, USA) for 2 h. Collected samples were then passed over a gravity-flow column (Bio-Rad) containing size exclusion Sephacryl of pore size 400- HR (Sigma Aldrich, St. Louis, MI, USA) to separate GPMVs from residual cell debris. For leakage assays, the thiol-reactive fluorescent probe, CellTracker Orange (Thermo Scientific, Rochester, NY, USA) was first applied to cells at 1:1000 dilution and incubated for 1 h at 37 ^o^C. The phospholipid content of unlabeled and labeled final material was measured by total phosphate assay.

### GPMV Imaging

Images were obtained in 8-well NUNC chambers (Thermo Scientific, Rochester, NY, USA) containing 250 μl of GMPV at 5 μM phospholipid lipid concentration. Imaging was carried out at the Yale Department of Molecular, Cellular, and Developmental Biology imaging facility, on a Zeiss LSM 510 confocal microscope, using a 63x Plan-Apo/1.4-NA oil-immersion objective with DIC capability (Carl Zeiss, Oberkochen, Germany). Image acquisition and processing were achieved using Zeiss Efficient Navigation (ZEN) and Image J software (Schneider et al., 2012).

### Fluorescence correlation spectroscopy (FCS)

FCS measurements were made on an LSM 880 Airyscan system NLO/FCS Confocal microscope (Carl Zeiss, Oberkochen, Germany) with a C-Apochromat 40×/1.2NA UV-VIS-IR Korr. water immersion objective (Carl Zeiss, Oberkochen, Germany). Thiol-conjugated molecules were excited at 594 nm. Confocal pinhole diameter was adjusted to 70 μm. Emission signals were detected through a 607-nm long-pass filter. Measurements were made in 8-well chambered coverglasses (Nunc, Rochester, NY, USA). All samples were incubated in GPMV buffer (10 mM HEPES, 150 mM NaCl, 2 mM CaCl_2_ (pH = 7.4)) for 1 h prior to taking measurements. Autocorrelation data were collected at regular intervals (5 min) with each autocorrelation curve collected over 10 s with 30 repeats.

Autocorrelations were fit using the software QuickFit3.0 (JW. Krieger, 2015). For thiol-conjugates, a model for a single diffusing species undergoing 3D Brownian diffusion was used.

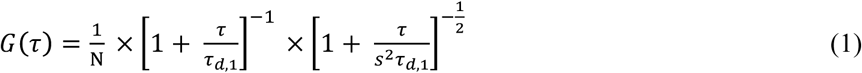

Here, *N* is the total thiol-conjugated molecules in the detection volume. The characteristic translational diffusion time of a diffusing particle is given by τ_*d*,1_. The structure factor, *s*, was determined as a free parameter for solutions of free Alexa 594 hydrazide dye and then fixed to the experimentally determined value of 0.1 for all subsequent fittings. For experiments of conjugates eluting during the second decay phase of leakage diffusion was assessed by burst counts within the time frame analysed.

### Confocal microscopy, cell imaging

Images were obtained in 8-well NUNC chambers (Thermo Scientific, Rochester, NY, USA) seeded with 20000 - 25000 cells/well. Cells were cultured for 48 h after passage before beginning experiments. For time dependent colocalization experiments of IAPP with JC-1 mitochondrial staining, the medium contained 200 nM IAPP_A647_, 13 μM unlabelled peptide. For experiments in the presence of ADM-116, 13 μM of molecule was introduced in the medium following a 3 h incubation of cells with IAPP. For experiments monitoring mitochondria, JC-1 was incubated with cells at 1:5000, at 37°C for 45 minutes prior to addition of protein. Images were acquired after 48 h total incubation time. Imaging was carried out using a x100 Plan-Apo/1.4-NA oil-immersion objective with DIC capability (Carl Zeiss, Oberkochen, Germany). For all experiments reporting on the colocalization of labelled IAPP, the gain setting for the blue channel was kept constant from sample to sample. Mitochondria containing JC-1 aggregates were detected in red (excitation 540 nm, emission 570 nm), and monomers in the green channel (excitation 490 nm, emission 520 nm). Image acquisition and processing were achieved using Zeiss Efficient Navigation (ZEN) and Image J software (Schneider et al., 2012).

### Intracellular imaging Förster resonance energy transfer (FRET)

The INS-1 growth media was replaced with media containing 100 nM IAPP_A488_ and 100 nM IAPP_A647_ (or 100 nM IAPP_A594_) and unlabelled IAPP and incubated for the time scales indicated in the text. Media was replaced with fresh prior to imaging. In experiments where ADM-116 was used, small molecule was added 3 h after incubation with protein. Images were acquired using a 100x Plan-Apo/1.4-NA oil-immersion objective with DIC capability (Carl Zeiss, Oberkochen, Germany). For the donor channel, IAPPA488 was excited with a 488 nm Argon2 laser and detected through a 505/550 nm emission filter. For the acceptor channel, IAPP_A647_ was excited with a 633 Argon2 laser and detected through a 730/750-nm emission filter. For all experiments the pinhole was kept constant to the Z-slice thickness of each filter channel. Single cell images were obtain for donor alone, acceptor alone and donor-acceptor fusion channels. Image acquisition and processing were achieved using Zeiss Efficient Navigation (ZEN) and Image J software (Soty M, 2011). The Image J plugin, RiFRET (Roszik et al., 2009), was used to calculate and remove the bleed through for each channel and to calculate a pixel-based FRET efficiency. The FRET distance was then calculated using:

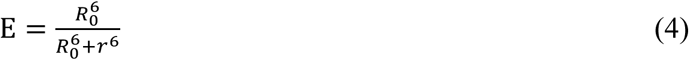

Where *E* is the calculated efficiency of FRET energy transfer, *R_0_* is the Förster distance and *r* is the distance between the donor and the acceptor.

### Imaging FRET

GPMV experiments were conducted in 8-well NUNC chambers (Thermo Scientific, Rochester, NY, USA) including 250 μM of GMPV stock solution at 5 μM apparent phospholipid (in monomer units). Wells containing GPMVs were mixed with 100 nM IAPP_A488_ and 100 nM IAPP_A647_ (or 100 nM IAPP_A594_) and unlabelled IAPP. In experiments where ADM-116 was used, small molecule was added at the same time as protein. The same microscope setup and analysis procedure was used to image FRET in GPMVs and cells.

### Cell culture

Rat insulinoma INS-1 cells (832/13, Dr. Gary W. Cline, Department of Internal Medicine, Yale University) were cultured at 37°C and 5% CO_2_ in phenol red free RPMI 1640 media supplemented with 10% fetal bovine serum, 1% penicillin/streptomycin (Life Technologies, Carlsbad, CA, USA), and 2% INS-1 stock solution (500 mM HEPES, 100 mM L-glutamine, 100 mM sodium pyruvate, and 2.5 mM β-mercaptoethanol). Cells were passaged upon reaching ~95% confluence (0.25% Trypsin-EDTA, Life Technologies), propagated, and/or used in experiments. Cells used in experiments were pelleted and resuspended in fresh media with no Trypsin-EDTA.

### Cell Viability

Cell viability was measured colorimetrically using the Cell-Titer Blue (CTB, Promega, Madison, WI, USA) fluorescence-based assay. Cells were plated at a density 5000 cells/well in 96-well plates (BD Biosciences, San Diego, CA). Peptide was directly introduced to each well after 48 h of culture and then incubated for an additional 48 h. For time dependent experiments, cells were incubated with peptide for the specified time points. After the incubation period, 20 μL CTB reagent was added to each well and incubated at 37°C and 5% CO_2_ for 2.5 – 3.5 h. Fluorescence of the resorufin product was measured on a FluoDia T70 fluorescence plate reader (Photon Technology International, Birmingham, NJ, USA). All wells included the same amount of water to account for different concentrations of peptide added to sample wells. Wells that included water vehicle but not peptide served as the negative control (0% toxic), and wells containing 10% DMSO were the positive control (100% toxic). Percent toxicity was calculated using the following equation:

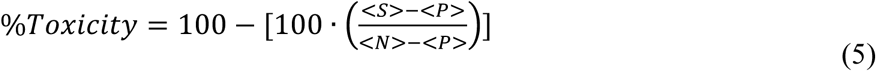

Each independent variable is the average of eight plate replicates from the negative control (*<N*>), positive control (<*P*>), and samples (<*S*>). Results presented for viability experiments are an average of three such experiments conducted independently on different days. Error bars represent the standard error of the mean.

### Phosphate assay

Lipid concentrations for GPMV preparations were determined by measuring total phosphate (King, 1932), assuming that all measured phosphate is from phospholipids, and that all lipids are phospholipids. This is a practical assumption designed to ensure reproducibility.

### Simulation of diluted-FRET

Expected values of FRET_e_ff for dynamic, isotropic distributions of spherically distributed IAPP were computed by numerical integration coded in-house using Mathematica (Wolfram Research, Champaign, IL). Briefly, the experimental ratio of labeled to unlabeled protein defines a binomial probability for the number of fluorophores distributed within trial spheres. The size of the spheres was dictated by constants (stated in the main text) for protein density and the total number of placed protein molecules. Spherical oligomers were generated randomly until the cumulative FRET_eff_ converged to a single value. Occurrences of multiple acceptors and donors within a single oligomer were accommodated by treating each potential dye pair, *ij,* as having an independent probability of resonance transfer, *p_ij_*. For example, a trial oligomer containing a single donor and three acceptors located a distance of R_o_ away gives a FRET_eff_ =0.875. Effects of donor-donor and acceptor-acceptor interactions were not considered. In general, multi-fluorophore corrections contribute negligibly to computations performed at the experimental ratios used in this work (Figure 7). The fraction of oligomers of size N with one donor and one acceptor at a doping ratio of 1:1:X is binomial with a probabilty of (2/(X+2)). This is further scaled by 0.5 to reflect the frequence that the two labelled IAPPs within the oligomer are donor and acceptor. Higher degrees of labelling within oligomers are rare at the doping ratios used in this work and so are ignored. To calculate the number of acceptor molecules contributing to the FRET signal detected, the number of pixels showing FRET was divided by the total number of acceptor pixels obtained from the acceptor channel.

## References

Aguzzi, A., and Altmeyer, M. (2016). Phase Separation: Linking Cellular Compartmentalization to Disease. Trends in cell biology 26, 547-558.

Antonenko, Y.N., Khailova, L.S., Knorre, D.A., Markova, O.V., Rokitskaya, T.I., Ilyasova, T.M., Severina, II, Kotova, E.A., Karavaeva, Y.E., Prikhodko, A.S., et al. (2013). Penetrating cations enhance uncoupling activity of anionic protonophores in mitochondria. PloS one 8, e61902.

Axelsen, P.H. (2008). A chaotic pore model of polypeptide antibiotic action. Biophysical journal 94, 1549-1550.

Brangwynne, C.P., Mitchison, T.J., and Hyman, A.A. (2011). Active liquid-like behavior of nucleoli determines their size and shape in Xenopus laevis oocytes. Proceedings of the National Academy of Sciences of the United States of America 108, 4334-4339.

Brender, J.R., Salamekh, S., and Ramamoorthy, A. (2012). Membrane disruption and early events in the aggregation of the diabetes related peptide IAPP from a molecular perspective. Accounts of chemical research 45, 454-462.

Costes, S., Langen, R., Gurlo, T., Matveyenko, A.V., and Butler, P.C. (2013). beta-Cell failure in type 2 diabetes: a case of asking too much of too few? Diabetes 62, 327-335.

Courchaine, E.M., Lu, A., and Neugebauer, K.M. (2016). Droplet organelles? The EMBO journal 35, 1603-1612.

Cremers, C.M., Knoefler, D., Gates, S., Martin, N., Dahl, J.U., Lempart, J., Xie, L., Chapman, M.R., Galvan, V., Southworth, D.R., et al. (2016). Polyphosphate: A Conserved Modifier of Amyloidogenic Processes. Molecular cell 63, 768-780.

Elbaum-Garfinkle, S., Ramlall, T., and Rhoades, E. (2010). The role of the lipid bilayer in tau aggregation. Biophys J 98, 2722-2730.

Gao, M., and Winter, R. (2015). The Effects of Lipid Membranes, Crowding and Osmolytes on the Aggregation, and Fibrillation Propensity of Human IAPP. Journal of diabetes research 2015, 849017.

Gregory, S.M., Cavenaugh, A., Journigan, V., Pokorny, A., and Almeida, P.F. (2008). A quantitative model for the all-or-none permeabilization of phospholipid vesicles by the antimicrobial peptide cecropin A. Biophysical journal 94, 1667-1680.

Gurlo, T., Ryazantsev, S., Huang, C.J., Yeh, M.W., Reber, H.A., Hines, O.J., O'Brien, T.D., Glabe, C.G., and Butler, P.C. (2010). Evidence for proteotoxicity in beta cells in type 2 diabetes: toxic islet amyloid polypeptide oligomers form intracellularly in the secretory pathway. The American journal of pathology 176, 861O–869.

Jayasinghe, S.A., and Langen, R. (2007). Membrane interaction of islet amyloid polypeptide. Biochimica et biophysica acta 1768, 2002–2009.

JW. Krieger, J.L. (2015). QuickFit 3.0: A data evaluation application for biophysics. In http://wwwdkfzde/Macromol/quickfit/.

Kegulian, N.C., Sankhagowit, S., Apostolidou, M., Jayasinghe, S.A., Malmstadt, N., Butler, P.C., and Langen, R. (2015). Membrane Curvature-sensing and Curvature-inducing Activity of Islet Amyloid Polypeptide and Its Implications for Membrane Disruption. The Journal of biological chemistry 290, 25782–25793.

King, E.J. (1932). The colorimetric determination of phosphorus. The Biochemical journal 26, 292–297.

Knowles, T.P., Vendruscolo, M., and Dobson, C.M. (2014). The amyloid state and its association with protein misfolding diseases. Nature reviews Molecular cell biology 15, 384–396.

Kumar, S., Birol, M., and Miranker, A.D. (2016a). Foldamer scaffolds suggest distinct structures are associated with alternative gains-of-function in a preamyloid toxin. Chemical communications 52, 6391–6394.

Kumar, S., Birol, M., Schlamadinger, D.E., Wojcik, S.P., Rhoades, E., and Miranker, A.D. (2016b). Foldamer-mediated manipulation of a pre-amyloid toxin. Nature communications 7, 11412.

Laganowsky, A., Liu, C., Sawaya, M.R., Whitelegge, J.P., Park, J., Zhao, M., Pensalfini, A., Soriaga, A.B., Landau, M., Teng, P.K., et al. (2012). Atomic view of a toxic amyloid small oligomer. Science 335, 1228–1231.

Last, N.B., Schlamadinger, D.E., and Miranker, A.D. (2013). A common landscape for membrane-active peptides. Protein science : a publication of the Protein Society 22, 870–882.

LeVine, H., 3rd (1993). Thioflavine T interaction with synthetic Alzheimer's disease beta-amyloid peptides: detection of amyloid aggregation in solution. Protein science : a publication of the Protein Society 2, 404–410.

Magzoub, M., and Miranker, A.D. (2012). Concentration-dependent transitions govern the subcellular localization of islet amyloid polypeptide. FASEB journal : official publication of the Federation of American Societies for Experimental Biology 26, 1228–1238.

Mukherjee A, M.-S.D., Butler PC, Soto C. ( 2015). Type 2 Diabetes as a Protein Misfolding Disease. Trends in molecular medicine 21, 439–449.

Mukherjee, A., Morales-Scheihing, D., Butler, P.C., and Soto, C. (2015). Type 2 diabetes as a protein misfolding disease. Trends Mol Med 21, 439–449.

Nath, A., Miranker, A.D., and Rhoades, E. (2011). A membrane-bound antiparallel dimer of rat islet amyloid polypeptide. Angewandte Chemie 50, 10859–10862.

Perry, R.J., Zhang, D., Zhang, X.M., Boyer, J.L., and Shulman, G.I. (2015). Controlled-release mitochondrial protonophore reverses diabetes and steatohepatitis in rats. Science 347, 1253–1256.

Podkalicka, J., Biernatowska, A., Majkowski, M., Grzybek, M., and Sikorski, A.F. (2015). MPP1 as a Factor Regulating Phase Separation in Giant Plasma Membrane-Derived Vesicles. Biophys J 108, 2201–2211.

Roszik, J., Lisboa, D., Szollosi, J., and Vereb, G. (2009). Evaluation of intensity-based ratiometric FRET in image cytometry--approaches and a software solution. Cytometry Part A : the journal of the International Society for Analytical Cytology 75, 761–767.

Schlamadinger, D.E., and Miranker, A.D. (2014). Fiber-dependent and -independent toxicity of islet amyloid polypeptide. Biophys J 107, 2559–2566.

Schneider, C.A., Rasband, W.S., and Eliceiri, K.W. (2012). NIH Image to ImageJ: 25 years of image analysis. Nature methods 9, 671–675.

Seeliger, J., Weise, K., Opitz, N., and Winter, R. (2012). The effect of Abeta on IAPP aggregation in the presence of an isolated beta-cell membrane. Journal of molecular biology 421, 348–363.

Sezgin, E., Kaiser, H.J., Baumgart, T., Schwille, P., Simons, K., and Levental, I. (2012). Elucidating membrane structure and protein behavior using giant plasma membrane vesicles. Nat Protoc 7, 1042–1051.

Soty M, V.M., Soriano S, del Carmen Carmona M, Nadal Á, Novials A. (2011). Involvement of ATP-sensitive Potassium (KATP) Channels in the Loss of Beta-cell Function Induced by Human Islet Amyloid Polypeptide. The Journal of biological chemistry 286, 40857–40866.

Su, X., Ditlev, J.A., Hui, E., Xing, W., Banjade, S., Okrut, J., King, D.S., Taunton, J., Rosen, M.K., and Vale, R.D. (2016). Phase separation of signaling molecules promotes T cell receptor signal transduction. Science 352, 595–599.

Williamson, J.A., Loria, J.P., and Miranker, A.D. (2009). Helix stabilization precedes aqueous and bilayer-catalyzed fiber formation in islet amyloid polypeptide. Journal of molecular biology 393, 383–396.

Wimley, W.C. (2010). Describing the mechanism of antimicrobial peptide action with the interfacial activity model. ACS chemical biology 5, 905–917.

